# Anxiety-like behavior during protracted morphine withdrawal is driven by gut microbial dysbiosis and attenuated with probiotic treatment

**DOI:** 10.1101/2025.01.29.633224

**Authors:** Mark Oppenheimer, Junyi Tao, Shamsudheen Moidunny, Sabita Roy

## Abstract

The development of anxiety during protracted opioid withdrawal heightens the risk of relapse into the cycle of addiction. Understanding the mechanisms driving anxiety during opioid withdrawal could facilitate the development of therapeutics to prevent negative affect and promote continued abstinence. Our lab has previously established the gut microbiome as a driver of various side effects of opioid use, including analgesic tolerance and somatic withdrawal symptoms. We therefore hypothesized that the gut microbiome contributes to the development of anxiety-like behavior during protracted opioid withdrawal.

In this study, we first established a mouse model of protracted morphine withdrawal, characterized by anxiety-like behavior and gut microbial dysbiosis. Next, we used fecal microbiota transplantation (FMT) to show that gut dysbiosis alone is sufficient to induce anxiety-like behavior. We further demonstrate that probiotic therapy during morphine withdrawal attenuates the onset of anxiety-like behavior, highlighting its therapeutic potential. Lastly, we examined transcriptional changes in the amygdala of morphine-withdrawn mice treated with probiotics to explore mechanisms by which the gut-brain axis mediates anxiety-like behavior.

Our results support the use of probiotics as a promising therapeutic strategy to prevent gut dysbiosis and associated anxiety during opioid withdrawal, with potential implications for improving treatment outcomes in opioid recovery programs.

## Introduction

Opioid use disorder (OUD) is a chronic relapsing condition characterized by compulsive opioid consumption and the development of a negative emotional state upon withdrawal from opioid use [1]. While individuals report various reasons for initial opioid use, the avoidance of opioid withdrawal is consistently cited as the primary driver for continued use [2]. Opioid withdrawal syndrome encompasses a range of aversive symptoms caused by the abrupt cessation of chronic opioid use.

Physical symptoms of opioid withdrawal syndrome including nausea, diarrhea, fever, and insomnia begin hours after the last dose of opioid and peak after around 72 h [3]. While physical symptoms usually subside within a week of the last dose, affective symptoms such as anxiety and depression can persist for weeks or months [4]. Importantly, the severity of these negative emotional states correlates strongly with heightened opioid cravings[5]. As a result, studies repeatedly associate heightened anxiety during opioid withdrawal with an increased risk of relapse [6–9]. Despite this, the underlying pathology of emergent anxiety behaviors during opioid withdrawal has not been sufficiently explained. Characterizing the mechanisms underlying these behaviors could facilitate the development of therapeutics aimed at alleviating the negative emotional symptoms of opioid withdrawal, ultimately reducing the risk of relapse.

Previous work from our lab has established the gut microbiome as a driver of several comorbidities associated with opioid use [10–13]. Chronic opioid use has been shown to induce distinct alterations to the composition of the gut microbiome, leading to a state of dysbiosis that can impact both the immune system and the brain [13]. Specifically, morphine administration has been linked to reduced alpha diversity in the gut microbiome, indicating suppressed microbial richness, and an increase in potentially pathogenic bacteria such as *E. faecalis* [14]. Furthermore, morphine compromises gut epithelial barrier integrity, enhancing the translocation of bacteria and their products, which can trigger pro-inflammatory immune responses [15]. Gut microbial dysbiosis induced by opioids has been linked to both the development of both morphine analgesic tolerance and somatic withdrawal symptoms. The development of morphine analgesic tolerance is attenuated in germ-free mice or by administration of antibiotics, implicating the gut microbiome as a contributor to this condition [16]. Administration of antibiotics has also been shown to reduce the somatic symptoms of morphine withdrawal, further establishing gut dysbiosis as a relevant factor in opioid use [17]. Based on this background, we hypothesized that the gut microbiome drives the development of anxiety-like behavior during protracted morphine withdrawal.

Collectively, clinical evidence and animal studies have clearly implicated the gut microbiome as a mediator of anxiety and depression [18, 19]. In humans, irritable bowel syndrome (IBS) is comorbid with anxiety disorders at rates as high as 30-50% [20]. Analysis of stool samples from patients with anxiety disorders reveals reduced microbial diversity and diminished populations of beneficial bacteria, such as Firmicutes and Tenericutes [21, 22]. Although clinical evidence on the efficacy of probiotics for anxiety and depression is limited [23], one study showed that probiotic supplementation alongside methadone maintenance treatment reduced depression severity in OUD patients [24].

Animal studies using germ-free mice, which have no commensal microbiota, have demonstrated altered anxiety-like behavior compared to conventional mice with an intact gut microbiome, further emphasizing the role of gut health in emotional regulation [25]. Studies varyingly report either elevated or reduced anxiety-like behavior in germ-free mice [26, 27], however the connection between the gut microbiome and anxiety is well-established.

Our understanding of negative affect associated with protracted opioid withdrawal has been significantly advanced through the use of animal models, particularly mice and rats [28]. Previous research has effectively modeled opioid withdrawal, demonstrating persistent negative affective behaviors such as anxiety and depression using an array of validated behavioral tests. Heightened anxiety-like and depression-like behavior has been reported in mice following one week of spontaneous morphine withdrawal [29, 30], lasting four- and seven weeks into protracted opioid withdrawal [31–33].

The establishment of these mouse models has enabled investigation of the neuropathology that drives the development of anxiety during opioid withdrawal. The amygdala, a brain region critical for emotional regulation, has consistently been implicated in this context [34–37]. Although the amygdala is highly innervated by serotonin and experimental manipulation of this signaling has been shown to alter anxiety-like behaviors [38], this signaling has not yet been implicated in the development of anxiety-like behavior following opioid withdrawal. Considering this background, the amygdala is an attractive target to manipulate therapeutically in effort to prevent the development of anxiety subsequent to opioid withdrawal.

In this study, we provided evidence that protracted morphine withdrawal is associated with gut microbial dysbiosis. Fecal microbiota transplantation from morphine-withdrawn mice to treatment-naive mice was sufficient to induce anxiety-like behavior. Additionally, probiotic therapy during protracted morphine withdrawal prevented anxiety-like behavior. Further, we propose altered serotonin signaling in the amygdala as a potential mechanism of the gut-brain axis that may mediate this behavior. These findings highlight probiotic therapy as a promising, clinically relevant adjunct to mitigate anxiety during opioid detoxification.

## Materials and Methods

### Animals

Female and male C57BL/6 mice between the ages of 12-15 weeks were purchased from Taconic Biosciences for use in experiments. Mice were housed in a facility which maintains a 12-hour light/dark cycle with constant temperature and humidity with unrestricted access to food and drinking water. All animal experiments were conducted in accordance with the guidelines of Institutional Animal Care and Use Committee (IACUC) at the University of Miami.

### Morphine treatment

Mice were subcutaneously implanted with a 75 mg slow-release morphine pellet (or inactive placebo) under isoflurane anesthesia. Briefly, an incision was made along the back of the animal where the pellet was inserted, and subsequently closed with surgical staples. Mice were removed from anesthesia and returned to home cage. After 72 h, surgical staples were removed and the same incision was reopened to facilitate removal of the pellet, inducing spontaneous withdrawal. As before, the incision was closed with surgical staples and mice were removed from anesthesia and returned to their home cage.

### Fecal microbiota transplantation

Before fecal microbiota transplantation (FMT), FMT recipient mice were first pre-treated with antibiotics to deplete their gut microbiome. FMT recipients received twice daily oral gavage of an antibiotic cocktail (neomycin: 100mg/kg, metronidazole: 100 mg/kg, vancomycin: 50 mg/kg) for 7 d. Additionally, home-cage drinking water was supplemented with 1 mg/ml ampicillin. FMT donor mice were subjected to protracted morphine withdrawal, or placebo treatment, as described above. On the eighth day following withdrawal, colon contents were harvested and processed for transplantation later the same day. Fresh colon contents from FMT donor mice were diluted in sterile PBS at a ratio of 1ml PBS/150mg and vortexed for one minute to homogenize. The mixture was filtered through a 75 μm cell strainer to separate out large solids. Approximately 36 h after the last dose of antibiotic cocktail, FMT recipient mice received 150 μl of the FMT solution by oral gavage. FMT recipient mice received a second dose of FMT six h after the first dose.

### Probiotic treatment

VSL#3 double strength powder was used for probiotic treatments. VSL#3 powder was suspended in nuclease free water at a ratio of 50mg VSL#3 per 150 µL water, yielding a concentration of approximately 1.6×10^9^ CFU per 150 µL dose, which was administered by oral gavage. Probiotic therapy was administered once daily at 5pm during protracted withdrawal, initiated the day of pellet removal and continued until the sixth day following pellet removal, for a total of six treatments.

### Behavioral testing

On the sixth day following pellet removal, mice completed the open field test as an assessment of anxiety-like behavior. Mice were moved from the housing facility to the testing room at 6pm and allowed to acclimate to the testing environment for two h before testing. The testing environment was kept under dark-light conditions with a white noise machine to cover background noise. Mice were tested using the Photobeam Activity System (PAS)-Open Field Test from San Diego Instruments, which employs a 16×16 photobeam floor grid to track activity. Mice placed in the center of the open field and allowed to roam freely for 30 minutes. Activity was recorded by the number of beam breaks, and activity in the center was defined by beam breaks in the center 4×4 of the grid. After the completion of testing, mice were returned to the housing facility.

On the seventh day following pellet removal, mice completed the elevated plus maze as a further assessment of anxiety-like behavior. Mice were moved to the testing environment at 6pm and allowed to acclimate to the testing environment for two h before testing. The testing environment was kept under red-light conditions with a white noise machine to cover background noise. Mice were placed in the center of the maze, facing an open arm, and allowed to explore the maze for 5 minutes. Activity was recorded by a camera positioned overhead which recorded video of the test. Videos were subject-blinded and manually scored. Time spent in the open and closed arms of the maze, as well as the number of entrances into the open and closed arms were recorded.

### Collection of biological specimens

The morning following behavioral testing, mice were sacrificed for collection of biological specimens. Mice were euthanized first by carbon dioxide chamber, followed by cervical dislocation. Amygdala samples were dissected from the brain according to Spijker, 2011 [39] and intestinal contents were harvested from the colon, all of which were flash frozen on dry ice for further analysis.

### DNA extraction and 16S rRNA gene sequencing

Intestinal contents were flash frozen on dry ice after collection and were stored at -80°C until ready for analysis. DNA was extracted from fecal samples using the DNeasy PowerSoil Pro kit (Qiagen; catalog no. 47016). Two extraction controls were included in sequencing to prevent contamination from the kit reagents. Sequencing was conducted by the University of Minnesota Genomics Center. The V4 region of the 16S rRNA gene was amplified by PCR using the forward primer 515F (GTGCCAFCMGCCGCGGTAA) and the reverse primer 806R (GGACTACHVGGGTWTCTAAT), along with Illumina adaptors and molecular barcodes, resulting in 427-bp amplicons. These amplicons were sequenced on the Illumina MiSeq version 3 platform, producing 300-bp paired-end reads.

### Microbiome analysis

Demultiplexed sequence reads were clustered into amplicon sequence variants (ASVs) with the DADA2 package (version 1.32.0) [40] in R (version 4.4.0) and RStudio (Build 764). The steps of the DADA2 pipeline include error filtering, trimming, learning of error rates, denoising, merging of paired reads, and removal of chimeras. ASVs were assigned taxonomically at the species level using a naive Bayesian classifier [41] in DADA2 with the SILVA reference database (release 138.1) [42]. The ASV and taxonomy tables were imported in MicrobiomeAnalyst [43] to create alpha and beta diversity plots, taxonomy bar plots, and linear discriminant analysis effect size (LEfSe) [44] plots. Low count and low variance ASVs were filtered with the default threshold. Total sum scaling was used for normalization. The threshold on the logarithmic LDA score for discriminative features was set to 2. The cutoff for *p*-value was set to 0.05 for LEfSe analysis. The Mann-Whitney test was used to detect if alpha diversity differed across treatments. Permutational multivariate analysis of variance (PERMANOVA) was used to detect if beta diversity differed across treatments.

### Transcriptome analysis

RNA sequencing samples was performed by Novogene Co. Raw data (raw reads) of fastq format were first processed through Novogene in-house perl scripts. In this step, clean data (clean reads) were obtained by removing reads containing adapter, reads containing poly-N and low-quality reads from raw data. At the same time, Q20, Q30 and GC content of the clean data were calculated. All the downstream analyses were based on clean data with high quality. Index of the reference genome was built using Hisat2 (v2.0.5) [45] and paired-end clean reads were aligned to the reference genome using Hisat2. FeatureCounts (v1.5.0) [46] from Subread package was used to count the reads numbers mapped to each gene. FPKM of each gene was calculated based on the length of the gene and reads count mapped to this gene. Differential expression analysis was performed using the DESeq2 [47] package (1.20.0). The resulting *p*-values were adjusted using the Benjamini and Hochberg approach for controlling false discovery rate. Genes with an adjusted *p*-value ≤ 0.05 found by DESeq2 were assigned as differentially expressed. Differentially expressed genes were subsequently analyzed using QIAGEN Ingenuity Pathway Analysis (IPA) for canonical pathways enrichment analysis. Fisher’s exact test was utilized in all those analyses to identify the signaling and metabolic pathways significantly associated with differentially expressed genes. Pathways with a *p*-value < 0.05 and a |z-score| >2 are considered significant.

### Statistical analysis

GraphPad Prism version 10.3.1 (GraphPad Software, Boston, Massachusetts USA, www.graphpad.com) was used for statistical analysis of behavioral data, as well as correlational- and linear regression modeling of transcriptome data. Details for statistical tests conducted for each experiment are found in figure legends. Outliers were identified using the ROUT method, with Q set to 1%.

## Results

### Protracted withdrawal from chronic morphine treatment is associated with heightened anxiety-like behavior in female and male mice

To model protracted morphine withdrawal, female (*n*=16-20 per group) and male (*n*=10-12 per group) mice were treated with a 75mg slow-release subcutaneous morphine pellet (or placebo) which was removed after 72 h to induce spontaneous withdrawal. On the sixth and seventh days following pellet removal, mice completed the open field test and elevated plus maze to assess anxiety-like behavior [Figure 1A].

**Figure 1.**
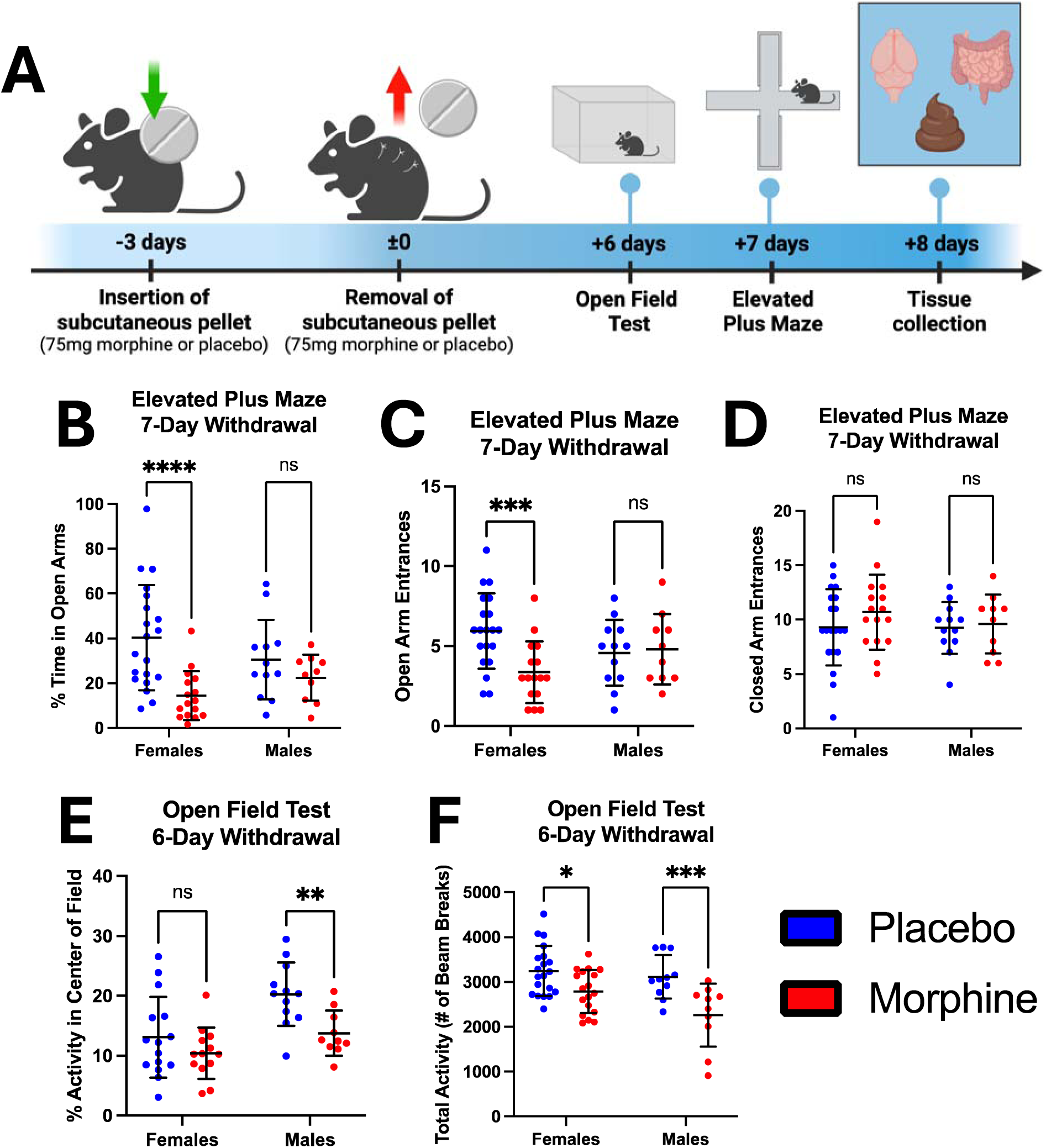
Protracted morphine withdrawal is associated with elevated anxiety-like behavior, with sex­ differences. (A) General experimental paradigm for morphine/placebo withdrawal and behavioral testing. (B) Percent time spent in the open arms of the elevated plus maze for female and male mice. (C) Number of entrances into the open arms of the elevated plus maze for female and male mice. (D) Number of entrances into the closed arms of the elevated plus maze for female and male mice. (E) Percent activity in the center of the open field test for female and male mice. (F) Total activity measured by number of beam breaks in the open field test for female and male mice. Symbols represent individual mice. ns= nonsignificant, *p<0.05, **p<0.01, ***p<0.001, ****p<0.0001 using Fisher’s Least Significant Difference test.

The elevated plus maze was the primary measure to assess anxiety-like behavior in both female and male mice withdrawn from chronic morphine treatment. Two-way ANOVA revealed a significant effect of treatment (F(1,54)=12.7, *p*=0.0008), but not sex (F(1,54)=0.0371, p=0.848) on time spent in the open arms of the maze. However, there was a nonsignificant trend towards an effect of the interaction of treatment and sex on time spent in the open arms of the maze (F(1,54)=3.48, *p*=0.0675). *Post hoc* analysis indicated that female mice withdrawn from chronic morphine treatment spent significantly less time in the open arms of the maze than female mice treated with placebo (t=4.40, *p*<0.0001) [Figure 1B]. In contrast, male mice withdrawn from chronic morphine treatment did not spend a significantly different percent of time in the open arms of the elevated plus maze than male mice treated with placebo (t=1.08, *p*=0.491) [Figure 1B]. Regarding the number of entrances into the open arms of the maze, two-way ANOVA indicated a significant effect of treatment (F(1,54)=4.04, *p*=0.0494), but not sex (F(1,54)=0.00247, *p*=0.961). In addition, the interaction of treatment and sex had a significant effect (F=5.66, *p*=0.0209) on the number of entrances into the open arms of the maze. According to *post hoc* analysis, female mice withdrawn from morphine made significantly fewer entrances into the open arms of the maze than did female mice treated with placebo (t=3.56, *p*=0.0016) [Figure 1C]. Male mice withdrawn from morphine treatment did not make a significantly different number of entrances into the open arms of maze than male mice treated with placebo (t=0.235, *p*=0.815) [Figure 1C]. There was no difference in the number of entrances into the closed arms of the maze between female mice withdrawn from morphine and placebo-treated females (t=1.31, *p*=0.197) [Figure 1D]. There was also no difference in the number of entrances into the closed arms of the maze between males withdrawn from morphine and placebo-treated males (t=0.258, *p*=0.797) [Figure 1D]. These results indicate induction of anxiety-like behavior in female mice, but not male mice, withdrawn from morphine treatment.

The open field test was utilized as a secondary measure of anxiety like behavior. Two-way ANOVA determined a significant effect of treatment on activity in the center of the field (F(1,46)=9.18, *p*=0.004), as well as a significant effect of sex (F(1,46)=11.9, *p*=0.0012). The interaction of sex and treatment had no significant effect on activity in the center of the open field (F(1,46)=1.58, *p*=0.215). *Post hoc* analysis demonstrated that male mice withdrawn from morphine exhibited significantly reduced activity in the center of the open field than placebo controls (t=2.86, p=0.0063) [Figure 1E]. In contrast, activity in the center of the open field was not significantly reduced in female mice withdrawn from morphine treatment, compared to placebo controls (t=1.34, *p*=0.187) [Figure 1E]. Regarding total activity in the open field test, there was a significant effect of treatment (F=19.1, *p*<0.0001) as well as a significant effect of sex (F=4.83, *p*=0.0322), but no significant effect of the interaction of treatment and sex (F(1,55)=1.76, *p*=0.190). *Post hoc* analysis reveals a significant decrease in total activity in the open field following protracted morphine withdrawal in both female (t=2.55, *p*=0.0134) and male mice (t=3.55, *p*=0.0008), compared to placebo controls [Figure 1E]. Results from the open field test indicate anxiety-like behavior in male mice withdrawn from morphine, and to a lesser extent suggest anxiety-like behavior in female mice withdrawn from morphine based on reduced total activity.

### Protracted withdrawal from chronic morphine treatment results in gut microbial dysbiosis in female and male mice

Previous studies from our lab have demonstrated gut dysbiosis associated with morphine dependence [14], and a partial recovery over the first 24 h of morphine withdrawal [17], however the state of the gut microbiome in the protracted stage of morphine withdrawal has not been described. To bridge this knowledge gap, we utilized 16S rRNA sequencing of colon contents collected from mice following behavioral testing. 16S rRNA sequencing demonstrated lasting alterations to the composition of the gut microbiome, consistent with gut microbial dysbiosis, in both female and male mice during protracted withdrawal from chronic morphine treatment. Beta diversity analysis, reported here by Bray-Curtis dissimilarity, revealed a significant shift in the composition of the gut microbiome following protracted morphine withdrawal in both female (F=2.61, *p*=0.014) [Figure 2A] and male mice (F=30.1, *p*=0.011) [Figure 2B]. More specifically, alpha diversity plotted by the Shannon index was increased in female mice withdrawn from morphine compared to placebo controls (U=97, *p*=0.0242) [Figure 2C], indicating an increase in species richness following protracted morphine withdrawal.

**Figure 2.**
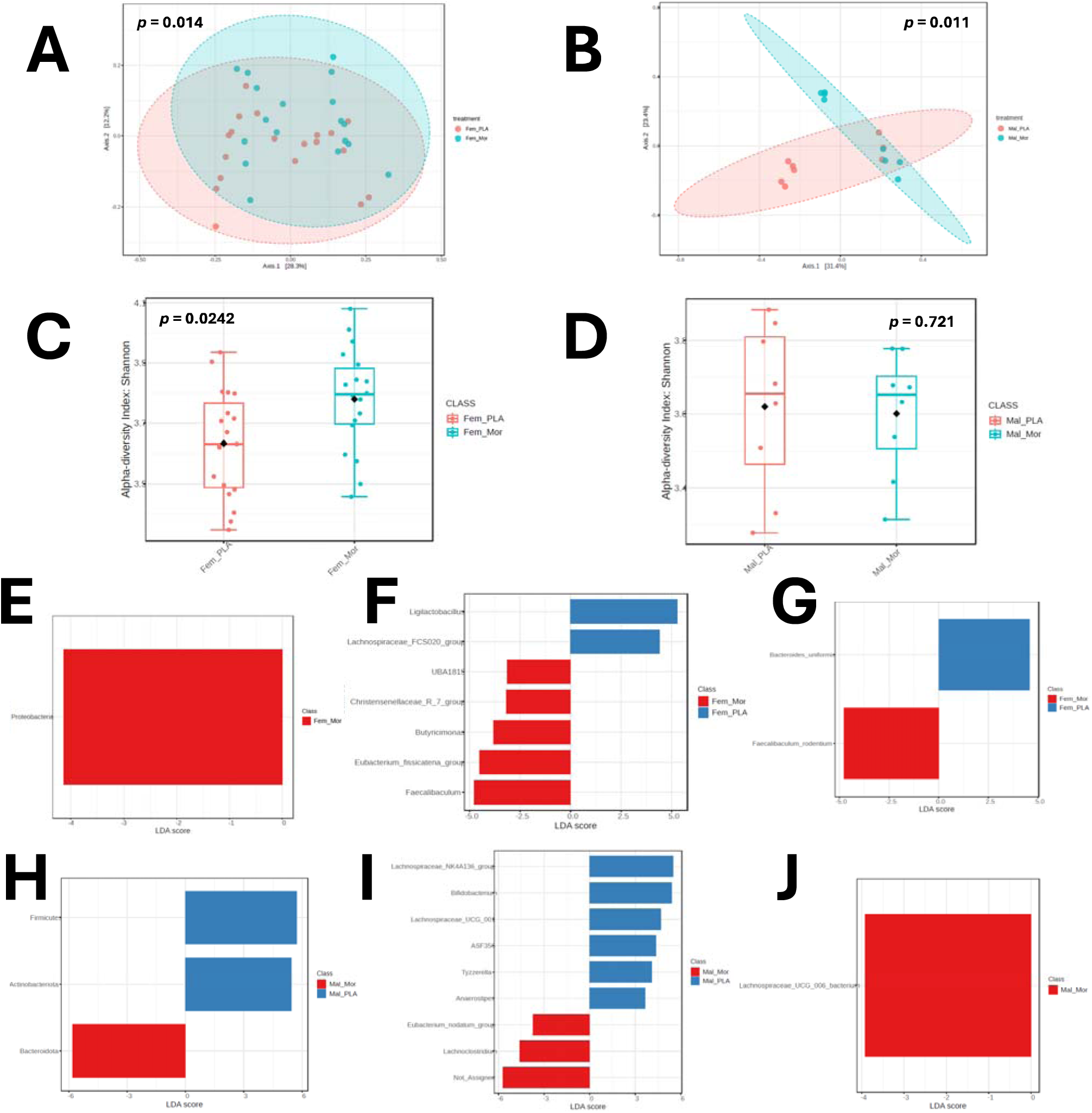
Gut microbial dysbiosis associated with protracted morphine withdrawal in female and male mice. (A) Principal coordinate analysis plot of Bray-Curtis distance (measure of -diversity) for female mice withdrawn from morphine and placebo controls. (B) Principal coordinate analysis plot of Bray-Curtis distance (measure of ­ diversity) for male mice withdrawn from morphine and placebo controls. (C) a-diversity plotted by Shannon index for female mice withdrawn from morphine and placebo controls. (D) a-diversity plotted by Shannon index for male mice withdrawn from morphine and placebo controls. (E) LEfSe plot for phyla enriched in morphine withdrawn­ female mice. (F) LEfSe plot for genera enriched and depleted in morphine-withdrawn female mice. (G) LEfSe plot for species enriched and depleted in morphine-withdrawn female mice. (H) LEfSe plot for phyla enriched and depleted in morphine-withdrawn male mice. (I) LEfSe plot for genera enriched and depleted in morphine-withdrawn male mice. (J) LEfSe plot for species enriched in morphine-withdrawn male mice. Fem_PLA = female placebo controls, Fem_Mor = female morphine withdrawal, Mal_PLA male placebo controls, Mal_Mor = male morphine withdrawal.

There was no such alteration in alpha diversity associated with morphine withdrawal in male mice (U=36, *p*=0.721) [Figure 2D]. This increase in alpha diversity seen in female mice withdrawn from morphine stands in contrast to the decrease in alpha diversity associated with morphine dependence [14], and represents an expansion during protracted withdrawal that overtakes placebo control levels.

Female mice withdrawn from morphine exhibited an enrichment of the phylum Proteobacteria compared to placebo controls [Figure 2E]. At the genus level, female mice withdrawn from morphine showed an expansion of *Faecalibaculum*, *Eubacterium fissicatena*, *Butyricimonas*, *Christensenellaceae R7*, and *UBA 1819* in addition to a depletion of *Ligilactobacillus* and *Lachnospiraceae FCS020* [Figure 2F]. At the species level, *Faecalibaculum rodentium* was expanded and *Bacteroides uniformis* was depleted in female mice withdrawn from morphine [Figure 2G].

Male mice withdrawn from morphine exhibited an enrichment of the phylum Bacteroidota, and depletion of the phyla Actinobacteria and Firmicutes compared to placebo controls [Figure 2H]. At the genus level, male mice withdrawn from morphine showed an expansion of *Lachnoclostridium*, *Eubacterium nodatum*, and an unassigned bacterial genus while the genera *Anaerostipes*, *Tyzzerella*, *ASF356*, *Lachnospiraceae UCG 001*, *Bifidobacterium*, and *Lachnospiraceae NK4A136* were depleted [Figure 2I]. The species *Lachnospiraceae UCG 006* bacterium was expanded in male mice withdrawn from morphine [Figure 2J].

Expansion of Proteobacteria coupled with the depletion of probiotic groups such as *Bacteroides uniformis* provided evidence of gut dysbiosis in female mice withdrawn from morphine. The altered ratio of Bacteroidota and Firmicutes abundance observed in male mice withdrawn from morphine, as well as depletion of probiotic flora such as *Bifidobacterium*, similarly indicated a dysbiotic gut microbiome.

### Fecal microbiota transplantation from morphine-withdrawn mice to treatment-naive mice induces anxiety-like behavior

After observing both gut dysbiosis and anxiety-like behavior following protracted morphine withdrawal, we set out to interrogate a causal relationship between these two adverse outcomes of morphine withdrawal. Specifically, we sought to understand if the state of gut dysbiosis associated with protracted morphine withdrawal was able to induce anxiety-like behavior in mice untreated by morphine. To accomplish this, we devised an experimental paradigm utilizing fecal microbiota transplantation (FMT) from female mice withdrawn from morphine, or placebo controls, to treatment-naive female mice [Figure 3A]. FMT donor mice completed behavioral testing during protracted morphine withdrawal (*n*=6), or placebo treatment (*n*=10), as previously described. Five mice from each group exhibiting the most significantly different behavior on the elevated plus maze [Figures 3B, 3C, 3D] were selected to have their gut microbiome transplanted to antibiotic-treated FMT recipients (*n*=10/group). One mouse in the Placebo FMT group and three mice in the Morphine FMT group made fewer than five combined entrances into either the open or closed arms of the elevated plus maze, and thus were excluded as outliers.

**Figure 3.**
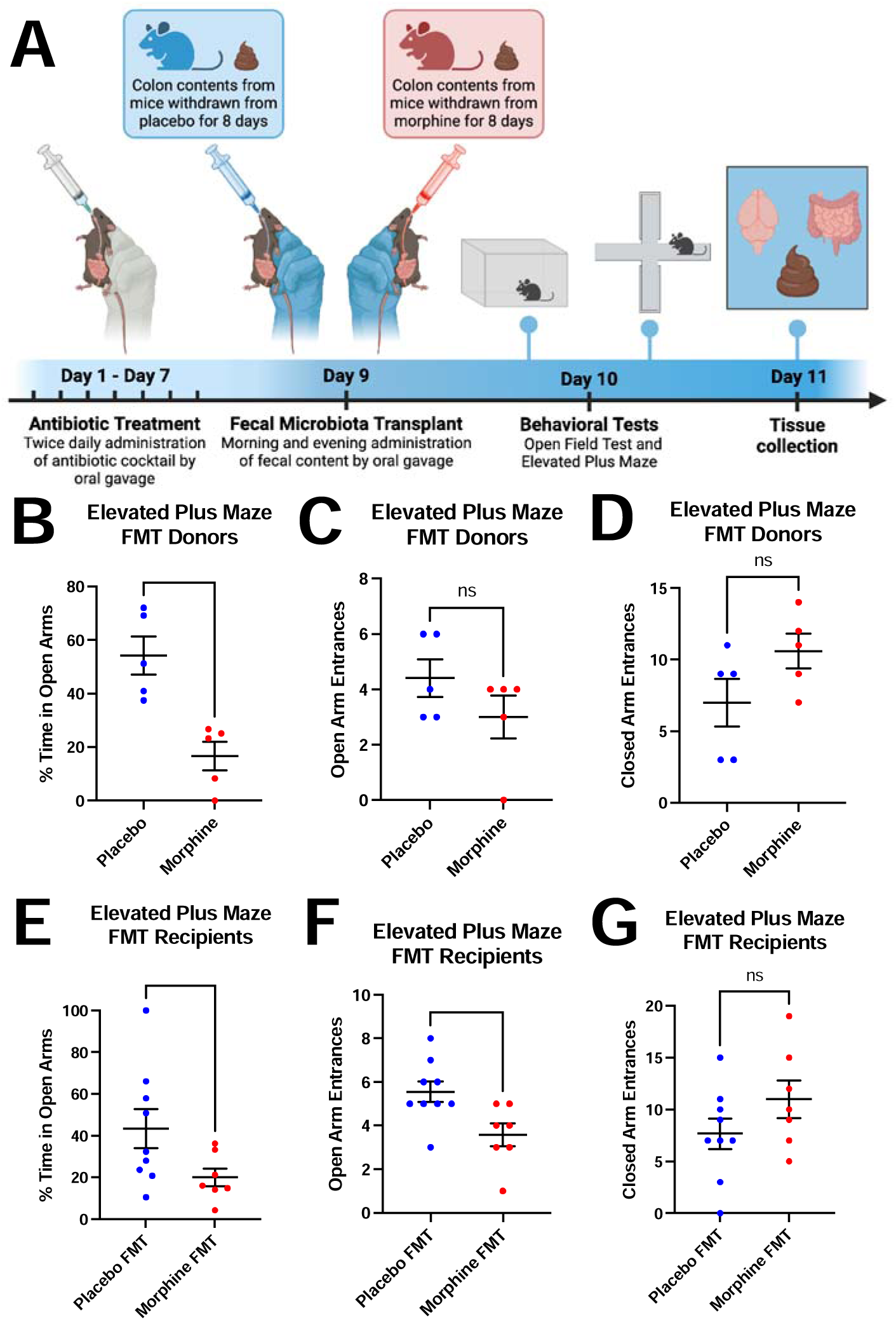
Fecal microbiota transplantation (FMT) from mice withdrawn from morphine results in increased anxiety-like behavior in the elevated plus maze. (A) Experimental paradigm for FMT and behavioral testing. (B) Percent time spent in the open arms of the elevated plus maze for FMT donor mice withdrawn from morphine and placebo controls. (C) Number of entrances into the open arms of the elevated plus maze for FMT donor mice withdrawn from morphine and placebo controls. (D) Number of entrances into the closed arms of the elevated plus maze for FMT donor mice withdrawn from morphine and placebo controls. (E) Percent time spent in the open arms of the elevated plus maze for mice who received FMT from morphine withdrawn mice and placebo controls. (F) Number of entrances into the open arms of the elevated plus maze for mice who received FMT from morphine withdrawn mice and placebo controls. (G) Number of entrances into the closed arms of the elevated plus maze for mice who received FMT from morphine withdrawn mice and placebo controls. ns= nonsignificant, *p<0.05, **p<0.01 using unpaired t-test with Welch’s correction

Mice that received FMT from donors withdrawn from morphine (Morphine FMT) spent significantly less time in the open arms of the elevated plus maze than mice that received FMT from donors treated with placebo (Placebo FMT) (t=2.264, *p*=0.0447) [Figure 3D]. Additionally, Morphine FMT mice made fewer entrances into the open arms of the maze than Placebo FMT mice (t=2.787, *p*=0.0145) [Figure 3F]. There were no significant behavioral differences found in the open field test [Supplemental Figure 1]. These results indicate that Morphine FMT mice show elevated anxiety-like behavior relative to the Placebo FMT group, suggesting that the effect of the gut microbiome associated with morphine withdrawal alone is sufficient to induce anxiety-like behavior.

### Administration of probiotics during morphine withdrawal rescues the induction of anxiety-like behavior in female mice

After establishing gut dysbiosis associated with protracted morphine withdrawal, we next investigated the gut microbiome as a therapeutic target to prevent the development of anxiety-like behavior subsequent to opioid withdrawal. Because morphine withdrawal impacted female mice more than male mice in terms of anxiety-like behavior, gut dysbiosis, and transcriptional alterations in the above experiments, it was decided to utilize female mice for the following experimental design. Based on the results of 16S sequencing of the gut microbiome, we decided to apply a probiotic therapy to counteract the expansion of dysbiotic flora and depletion of probiotic flora observed in female mice withdrawn from morphine treatment. To this end, female mice (*n*= 10-12 per group) underwent spontaneous withdrawal as previously described, with the addition of once-daily administration of VSL#3 probiotic blend (or water control) by oral gavage throughout the protracted withdrawal phase until behavioral testing [Figure 4A].

**Figure 4.**
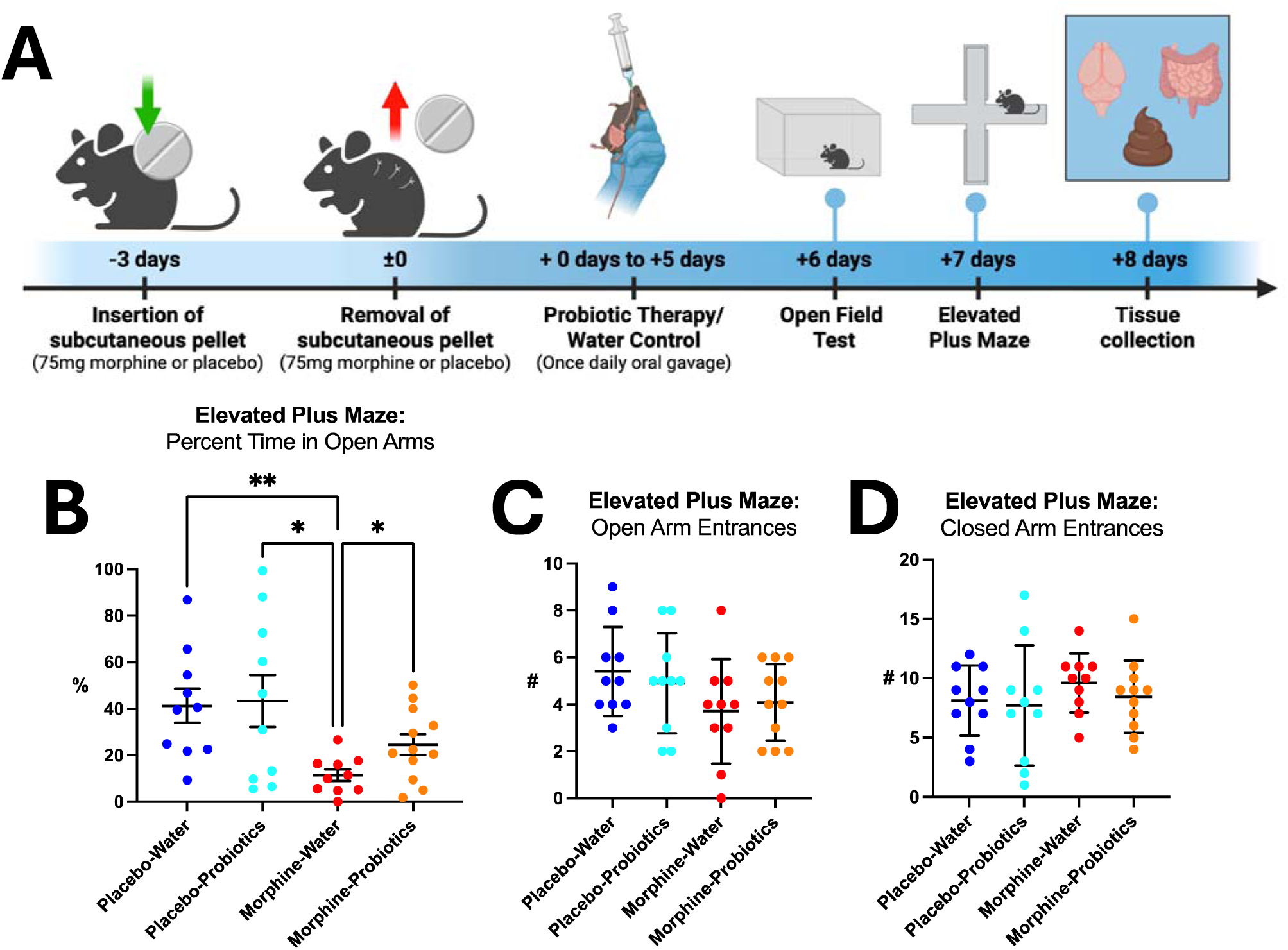
Probiotic therapy during protracted withdrawal partially rescues the development of anxiety-like behavior during protracted morphine withdrawal. (A) Experimental paradigm for probiotic therapy during protracted morphine withdrawal and behavioral testing. (B) Percent time spent in the open arms of the elevates plus maze. (C) Number of entrances into the open arms of the elevated plus maze. **(D)** Number of entrances into the closed arms of the elevated plus maze. Symbols represent individual mice; line and error bars represent mean and standard deviation. *p<0.05, **p<0.01 using unpaired t-test with Welch’s correction.

Comparing time spent in the open arms of the elevated plus maze, Welch’s ANOVA test determined a significant difference between the means of the four treatment groups (F(3,18.8)=7.53, *p=*0.0017). According to *post hoc* analysis, the morphine-water group spent significantly less time in the open arms of the maze than the placebo-water group (t=3.83, *p*=0.0028) [Figure 4B], confirming the induction of anxiety-like behavior in the positive control group. Importantly, the morphine-withdrawn females that received probiotic therapy spent significantly more time in the open arms of the maze than the morphine-water group (t=2.59, *p*=0.0191) [Figure 4B]. This indicates lower anxiety-like behavior in morphine treated females following probiotic therapy. Of note, there was no significant difference in time spent in the open arms of the maze between the morphine-probiotics group and the placebo-probiotics group (t=1.56, *p*=0.146) [Figure 4B]. One-way ANOVA analysis revealed no significant difference among the mean number of entrances into the open arms of the maze between any of the four treatment groups (F(3,37)=1.53, *p*=0.222) [Figure 4C]. Additionally, one-way ANOVA showed no significant difference in entrances into the closed arms of the maze among the four groups (F(3,37)=0.540, *p*=0.658) [Figure 4D]. These results show a rescue of the induction of anxiety-like behavior associated with protracted morphine withdrawal in female mice treated with VSL#3 probiotic blend.

In the open field test, there were no significant differences in levels of activity in the center of the field or total activity levels between any of the four treatment groups [Supplementary Figure 2A, 2B]. This was expected based on previous experiments using female mice.

Results from the elevated plus maze, our primary assessment of anxiety-like behavior, indicate that probiotic therapy during protracted withdrawal from chronic morphine treatment rescues the induction of anxiety-like behavior, implicating the gut microbiome as a driver of this behavior.

### Protracted morphine withdrawal induces transcriptomic changes in the amygdala of female and male mice

The gut microbiome can have significant impacts on the brain and behavior, and the amygdala has been identified as a critical signaling node across the gut-brain axis [48]. We endeavored to investigate potential mechanisms that mediate the impact of morphine withdrawal-induced gut dysbiosis on anxiety-like behavior by assessing the transcriptomic state of the amygdala during protracted withdrawal. To assess transcriptional alterations in the brain associated with protracted morphine withdrawal, amygdala samples collected from female and male mice (*n* = 6 per group) in the above paradigm of withdrawal and behavioral testing [Figure 1A] were submitted for bulk RNA-sequencing. Analysis of differential expressed genes (DEGs) between morphine-withdrawn and placebo-treated mice reveals a greater degree of transcriptional alteration in female mice than male mice [Figure 5A]. 2640 genes were identified as differentially expressed in female mice withdrawn from morphine compared to placebo controls, whereas 1154 DEGs were identified in morphine-withdrawn male mice compared to placebo controls. Overall, the pattern of differential expression favored a greater degree of upregulated-than downregulated genes for both female and male mice. Interestingly, this pattern resembles the results from behavioral testing which indicate more significant anxiety-like behavior in female mice withdrawn from morphine compared to male mice.

**Figure 5.**
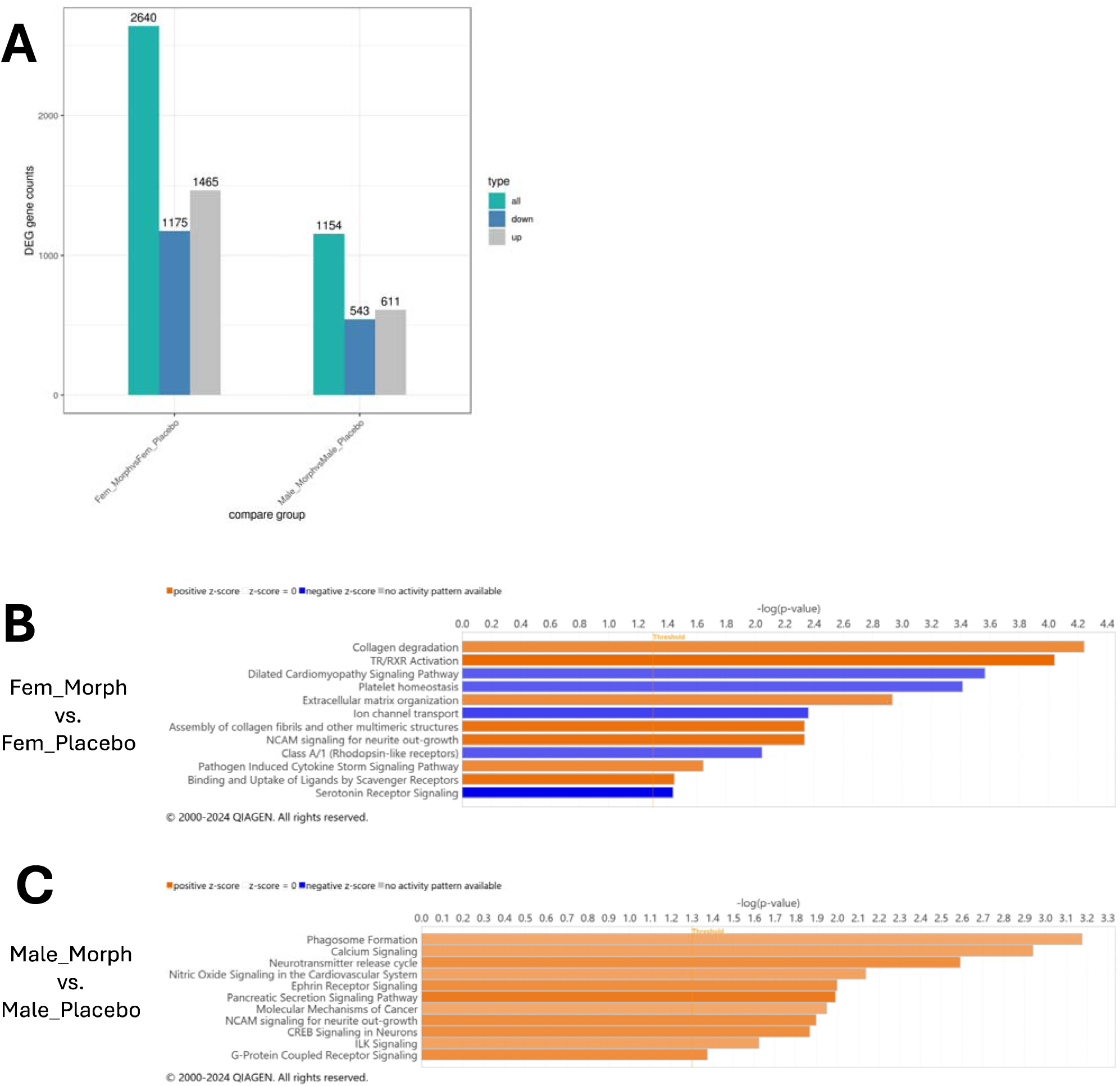
Transcriptional alterations to the amygdala associated with protracted morphine withdrawal in female and male mice. (A) Differentially expressed gene (DEG) counts for Fem_Morph vs. Fem_Placebo and Male_Morph vs. Male_Placebo. (B) Pathways identified by IPA as upregulated and downregulated in Fem_Morph vs. Fem_Placebo. (C) Pathways identified by IPA as upregulated in Male_Morph vs. Male_Placebo. Fem_Morph = female morphine withdrawal, Fem_Placebo = female placebo controls, Male_Morph male morphine withdrawal, Male_Placebo = male placebo controls.

Results from bulk RNA-sequencing were further contextualized using QIAGEN Ingenuity Pathway Analysis (IPA) software. IPA was used to examine patterns of gene expression and reveal canonical pathways that are predicted to be activated or inhibited in the context of protracted morphine withdrawal in female and male mice.

In female mice withdrawn from chronic morphine treatment, a number of pathways were identified as both significantly activated and inhibited [Figure 5B]. Of particular interest was the inhibition of serotonin receptor signaling, due to the well-established connection between serotonin and anxiety [49]. On this signaling pathway, genes downregulated in females withdrawn from morphine withdrawal included genes encoding multiple serotonin receptors, such as *Htr2c* and *Htr5b*, as well as the adrenergic receptor *Adra2a*. Interestingly, despite the more significant pattern of gene downregulation on the serotonin receptor signaling pathway, upregulation of several genes associated with this pathway was observed. These upregulated genes included *Htr6* and *Htr1d* which also encode for serotonin receptors, along with the adrenoreceptor *Adra2b*. IPA additionally revealed an upregulation of the pathogen-induced cytokine storm signaling pathway in morphine-withdrawn females, highlighting the immunomodulatory effects of protracted withdrawal. The identification of these pathways provides insight into potential mechanisms that regulate the development of anxiety-like behavior subsequent to morphine withdrawal.

Among male mice withdrawn from morphine, IPA showed only significant activated pathways with no significantly inhibited signaling pathways [Figure 5C]. Included in these signaling pathways was activation of G-protein coupled receptor signaling. On this pathway, upregulation of the genes encoding for adrenoceptor alpha 1B (*Adra1b*) as well as dopamine receptor D4 (*Drd4*) was observed, as well as downregulation of *Htr7* which encodes for a serotonin receptor. While sequencing results from male mice presented some individual genes of interest, the overall patterns of expression lacked the cohesiveness that characterized the sequencing data from female mice.

### Probiotic treatment during protracted morphine withdrawal is associated with transcriptomic changes in the amygdala of female mice

To explore potential mechanisms by which probiotic treatment during protracted morphine withdrawal was able to reduce the development of anxiety-like behavior in female mice, amygdala samples collected as part of the above experimental paradigm (*n*=3-6) were submitted for bulk RNA-sequencing. Consistent with our previous findings, DEG analysis revealed that female mice withdrawn from morphine and treated with water during withdrawal (MOR_Water) exhibited a higher degree of upregulated-than downregulated genes compared to placebo-withdrawn mice treated with water (PLA_Water) [Figure 6A]. Indeed, Ingenuity Pathway Analysis (IPA) revealed only significantly upregulated signaling pathways associated with the MOR_Water condition compared to PLA_Water [Figure 6B]. Interestingly, morphine-withdrawn mice treated with probiotics (MOR_Prob) showed a greater degree of downregulated genes compared to the MOR_Water condition [Figure 5A].

**Figure 6.**
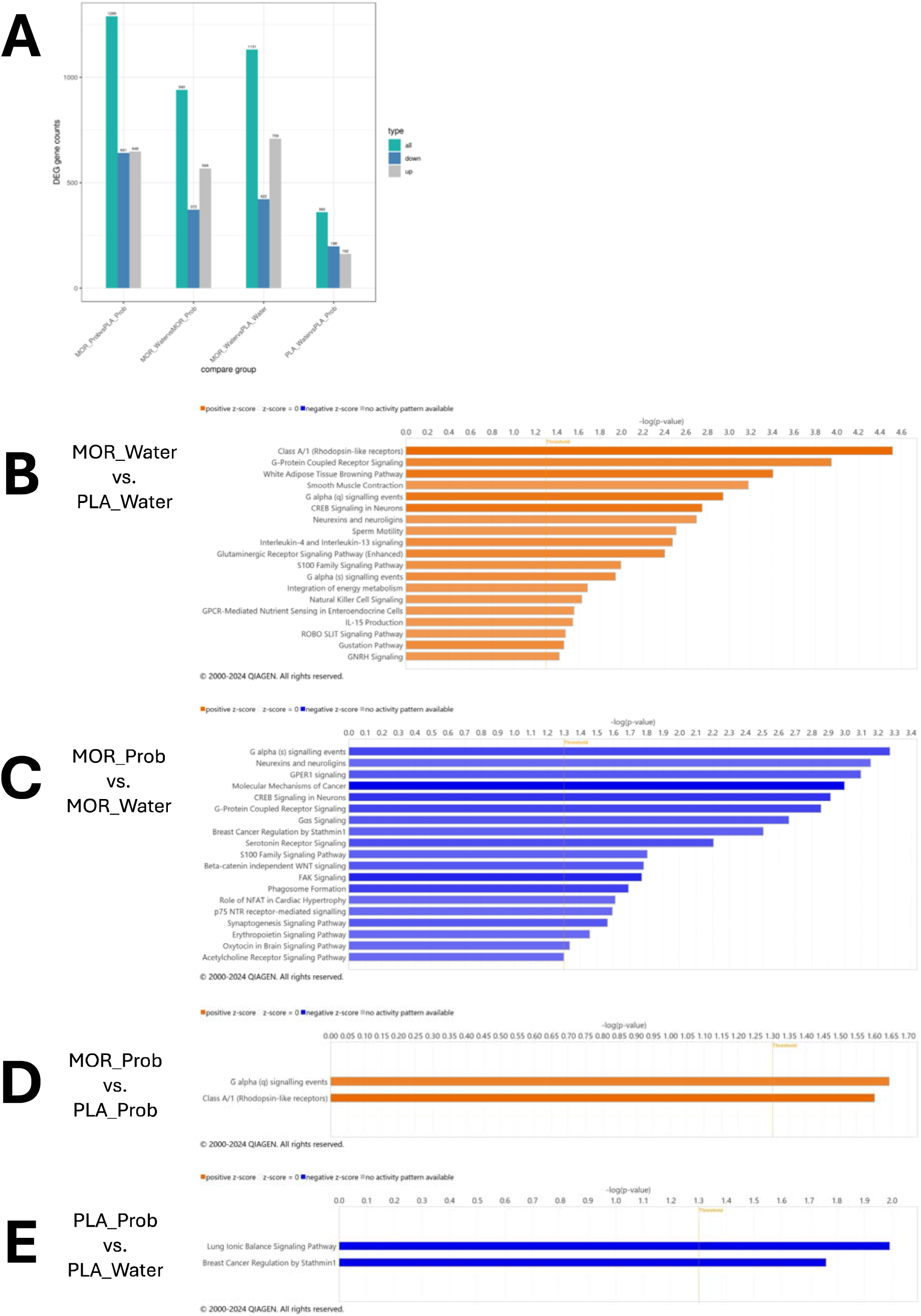
Transcriptional alterations to the amygdala associated with protracted morphine withdrawal and probiotic therapy. (A) Differentially expressed gene (DEG) counts for MOR_Prob vs. PLA_Prob, MOR_Water vs. MOR_Prob, MOR_Water vs. PLA_Water, and PLA_Water vs. PLA_Prob. (B) Pathways identified by IPA as upregulated in MOR_Water vs. PLA_Water. (C) Pathways identified by IPA as downregulated in MOR_Prob vs. MOR_Water. (D) Pathways identified by IPA as upregulated in MOR_Prob vs. PLA_Prob. (E) Pathways identified by IPA as downregulated in PLA_Prob vs. PLA_Water. MOR_Prob = morphine withdrawal, probiotic therapy. PLA_Prob = placebo treatment, probiotic therapy. MOR_Water = morphine withdrawal, water control. PLA_Water = placebo control, water control.

Accordingly, IPA showed only inhibited signaling pathways in the MOR_Prob condition compared to the MOR_Water condition [Figure 6C]. The implication of these results is that morphine withdrawal broadly upregulates multiple signaling pathways that are in turn downregulated by probiotic therapy. Probiotic treatment in the placebo condition (PLA_Prob) was associated with relatively minor differential gene expression compared to the PLA_Water condition [Figure 6A], suggesting that the transcriptional alterations induced by probiotic treatment were specific to morphine withdrawal.

Several pathways identified by IPA as significantly inhibited in the morphine-probiotics condition compared to the morphine-water condition were noted as potentially relevant to the development of anxiety-like behavior, including Gαs signaling, GPER1 signaling, and serotonin receptor signaling. Gαs signaling was inhibited following probiotic treatment during morphine withdrawal due to downregulation of the genes *Adcy5*, *Crhr1*, *Gng7*, *Htr6*, and *Mc3r*. Probiotic treatment. GPER1 signaling was similarly inhibited due to downregulation of *Adcy5*, *Cit*, *Gng7*, *Pik3c2b*, and *Stk32a*. Counterintuitively, probiotic treatment during morphine withdrawal was also associated with inhibition of serotonin receptor signaling. As part of this pathway, significant downregulation of a total of nine genes was observed: *Htr6*, *Adcy5*, *Cacna1i*, *Pik3c2b*, *Rasd2*, *Shank3*, *Htr1b*, *Gng7*, and *Rhobtb2*.

Based on the established connection between serotonin receptor signaling and anxiety, we further examined the expression of genes on this pathway that are differentially regulated by probiotic treatment during protracted morphine withdrawal. A correlation matrix was constructed to compare the expression of these genes with the time spent in the open arms of the elevated plus maze by the corresponding animal [Figure 7A]. Spearman correlation analysis identified three genes that were significantly negatively correlated with time spent in the open arms of the maze: *Htr6* (r=-0.797, *p*=0.003), *Pik3c2b* (r=-0.713, *p*=0.012), and *Rhobtb2* (r=-0.720, *p*=0.011). Additionally, a multiple linear regression was employed to assess how expression of the nine downregulated genes on the serotonin signaling pathway influence time spent in the open arms of the elevated plus maze. The overall fit of the model (R^2^= 0.993) was statistically significant (F(9,2)=33.43, *p*=0.0294), indicating that the model explains a significant portion of the variance in time spent in the open arms of the elevated plus maze [Figure 7B]. Fascinatingly, six of these nine downregulated genes were identified as significantly upregulated in morphine-withdrawn females compared to placebo controls in our RNA-sequencing study [Figures 7C-7H]. This internal consistency further strengthens the observation of altered serotonin signaling associated with protracted morphine withdrawal, and the amelioration of these signaling alterations following probiotic therapy suggests a mechanism by which the gut microbiome may function across the gut-brain axis to alleviate the development of anxiety-like behavior.

**Figure 7.**
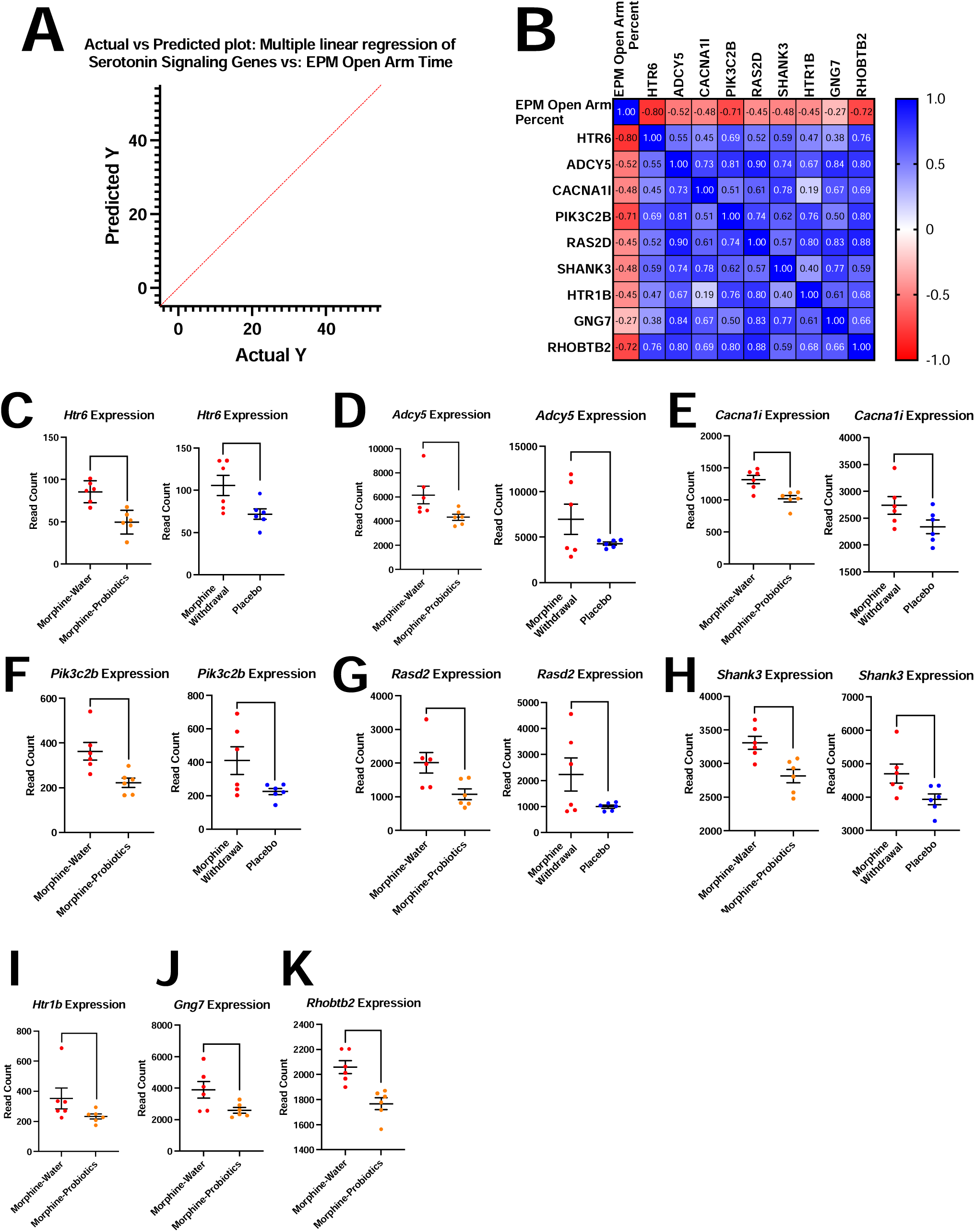
Altered expression of serotonin signaling genes associated with morphine withdrawal and probiotic therapy. (A) Multiple linear regression model: altered serotonin signaling genes vs. percent time spent in the open arms of the elevated plus maze. (B) Correlation matrix of serotonin signaling genes downregulated in morphine-probiotics vs. morphine-water and percent time spent in the open arms of the elevated plus maze. (C-H) Comparative alterations to serotonergic gene expression between morphine-water vs. morphine-probiotics and morphine vs. placebo females. (I-K) Serotonergic genes differentially expressed in morphine-water vs. morphine-probiotics, but not morphine vs. placebo females. Symbols represent individual mice; line and error bars represent mean and standard deviation. *p<0.05, **p<0.01, ***p<0.001 using the Wald test.

## Discussion

The development of anxiety symptoms is a common feature of protracted opioid withdrawal; however previous studies have not investigated the role of the gut microbiome in this behavior. In this study, we first show that protracted morphine withdrawal is associated with gut microbial dysbiosis.

Second, we provide evidence that morphine withdrawal-induced gut dysbiosis contributes to the development of anxiety-like behavior, which can be prevented by the administration of probiotics during withdrawal. Further, we propose a mechanism of altered amygdala serotonin receptor signaling that may mediate the interaction of the opioid-induced gut dysbiosis and anxiety-like behavior. These results advance our understanding of the gut microbiome during opioid withdrawal and consequently support the use of probiotic therapy for individuals discontinuing chronic opioid use.

Bulk-RNA sequencing data indicate gut dysbiosis in both female and male mice which persists into the protracted phase of morphine withdrawal. Specifically, there were significant shifts in beta diversity, indicating distinct differences in microbial composition between morphine-withdrawn mice and placebo controls. Interestingly, female mice demonstrated an increase in alpha diversity, or species richness, contrasting with the decrease in alpha diversity previously associated with morphine dependence [14]. This suggests the partial rebound of microbial richness observed during the first 24 h of morphine withdrawal [17] continued beyond the point of placebo controls over the following week. In both female and male mice, we showed an expansion of gut flora associated with inflammation and dysbiosis. In female mice, we observed an expansion of the phylum Proteobacteria, a proposed microbial signature of gut dysbiosis [50] which is linked to inflammatory bowel disease (IBD) [51, 52] and chronic stress [53]. Male mice withdrawn from chronic morphine treatment presented with an enrichment of Bacteroidota and a depletion of Firmicutes at the phylum level. A disturbance to the ratio of the abundance of these two phyla (F/B ratio) is indicative of gut dysbiosis [54]. Specifically, the decreased F/B ratio observed here in males withdrawn from morphine is typically associated with inflammatory bowel disease [54–56]. Both female and male mice also show a depletion of probiotic flora following morphine withdrawal. Female mice withdrawn from morphine showed a depletion of the *Ligilactobacillus* genus and the species, *Bacteroides uniformis*. *Bacteroides uniformis*, a commercially available probiotic, is associated with metabolic and immune benefits [57, 58] and has even been termed a “psychobiotic” due to its role in serotonin metabolism and reduction of anxiety-like behaviors [59]. Male mice withdrawn from morphine showed a notable depletion of Bifidobacterium, a genus widely recognized for its probiotic properties [60]. Bifidobacteria inhibit LPS-induced NF-κB activation [61], prevent pathogen expansion [62], and have been established to reduce anxiety in both mice [63] and humans [64]. Overall, protracted morphine withdrawal is associated with an expansion of dysbiotic gut flora along with a depletion of beneficial bacterial groups that are associated with anxiety like behavior.

Using fecal microbiota transplantation (FMT), we demonstrated that gut dysbiosis associated with morphine withdrawal is sufficient to induce anxiety-like behavior in treatment-naive mice. This finding supports a causal role for the gut microbiome in the development of negative affective, such as anxiety-like behavior, during morphine withdrawal. FMT includes the transfer of bacteria as well as their metabolites [65], which may also impact behavior [66]. Bacterial metabolites include bidirectional modulators of neuroinflammation, neuroactive mediators, neurotransmitter receptor agonists, and other products capable of orchestrating a speedy behavioral response [67]. Given the short treatment duration and relatively brief time before behavioral assessment employed in this experiment, the effect of the FMT we observed is likely a result of bacterial metabolites in the FMT solution rather than the eventual engraftment of bacterial groups. Future studies will investigate the metabolome associated with protracted morphine withdrawal to assess mechanisms of the gut-brain axis.

Building on our observation of depleted probiotic bacterial groups associated with morphine withdrawal, we investigated whether probiotic supplementation during protracted morphine withdrawal could mitigate the induction of anxiety-like behavior. To enhance clinical relevance, we selected a commercially available probiotic blend (VSL#3) for testing. Morphine-withdrawn mice treated with VSL#3 exhibited reduced anxiety-like behavior compared to morphine-withdrawn mice given a water gavage, suggesting an anxiolytic effect of the probiotic treatment. Probiotic therapy had no effect on placebo-treated mice, indicating that the anxiolytic effect was specific to morphine withdrawal. We propose that VSL#3 treatment countered the dysbiotic expansion observed via 16S sequencing by supplementing the depleted probiotic groups, potentially preventing the onset of anxiety-like behavior. These results support the feasibility of probiotic therapy as a potential adjunct treatment in opioid detox settings to alleviate some of the negative emotional effects of opioid withdrawal.

To identify mechanisms by which gut dysbiosis following protracted morphine withdrawal may contribute to anxiety-like behavior, we conducted bulk RNA-sequencing on amygdala samples from mice withdrawn from either morphine or placebo. The amygdala is known to influence hedonic processing during protracted morphine withdrawal [68] and to modulate the development of withdrawal-associated anxiety [34, 69]. Ingenuity Pathway Analysis (IPA) of the RNA-sequencing data revealed significant downregulation of serotonin signaling in morphine-withdrawn female mice, suggesting a possible link to anxiety-like behavior. Within this pathway, serotonin receptors *Htr2c* and *Htr5b* were notably upregulated. Prior studies indicate that *Htr2c* knockout can reduce anxiety-like behavior by dampening amygdala signaling [70], highlighting a potential role of *Htr2c* upregulation in anxiety during withdrawal. Interestingly, despite the overall downregulation of serotonin signaling, several pathway genes, including *Htr6*, were paradoxically upregulated.

To further investigate how the gut microbiome may impact anxiety-like behavior during morphine withdrawal, we conducted additional bulk RNA-sequencing on amygdala samples, this time collected from mice under the probiotic therapy paradigm. IPA showed a trend of predominantly upregulated pathways in the morphine withdrawal condition without probiotics, whereas probiotic treatment led to a generalized downregulation of these pathways. Many of these pathways downregulated by probiotic treatment during morphine withdrawal involved G-protein signaling, including downregulation of Gαs signaling. Conversely, activation of Gαs signaling in the amygdala has been shown to induce anxiety-like behavior [71, 72]. Interestingly, GPER1 (G protein-coupled estrogen receptor 1) signaling was also downregulated by probiotic treatment during morphine withdrawal. GPER1 binds and carries out effects of estradiol, the female sex hormone, and high serum levels of GPER1 have been associated with heightened anxiety in patients [73]. Accordingly, genetic knockout of GPER1 has been shown to reduce anxiety in female mice [74]. Most notably, serotonin signaling was downregulated in the morphine-probiotic condition compared to the morphine-water control. While this might seem counterintuitive, it is contextualized by examination of the genes that drive the expression of this pathway. For instance, *Htr1b* expression was reduced in the morphine-probiotic condition relative to morphine-water, aligning with evidence that 5-HT1B receptor antagonists can have antidepressive effects, while agonists may induce depressive-like behavior [75, 76]. Expression of *Htr6* was also lower in the morphine-probiotic condition, mirroring previous RNA-sequencing results that found elevated *Htr6* expression associated with morphine withdrawal. Intriguingly, both agonists and antagonists of the 5-HT6 receptor have shown anxiolytic effects [77], emphasizing the complex interplay between serotonin signaling and anxiety-related behaviors.

Of the nine serotonin signaling genes downregulated in the morphine-probiotic group, six had been upregulated in our initial RNA-sequencing analysis of morphine-withdrawn females (*Htr6*, *Adcy5*, *Cacna1i*, *Pik3c2b*, *Rasd2*, and *Shank3*). Expression of these genes also strongly predicted anxiety-like behavior in the elevated plus maze, as confirmed by linear regression analysis. Together, these findings suggest that morphine withdrawal upregulates specific serotonergic genes in female mice, which are subsequently downregulated following probiotic therapy. This pattern highlights a potential mechanism by which probiotic therapy may modulate the gut-brain axis to prevent anxiety-like behavior during morphine withdrawal.

These findings provide compelling evidence that protracted morphine withdrawal induces gut microbial dysbiosis, which in turn contributes to the development of anxiety-like behavior. By combining 16S rRNA sequencing, fecal microbiota transplantation, and probiotic therapy, we demonstrate both the causality of gut dysbiosis in the onset of anxiety-like behaviors and the potential of probiotic intervention to mitigate these effects. Our results further suggest a complex interaction between gut microbiota and central serotonin signaling in the amygdala, revealing alterations in serotonergic pathways that may underlie the behavioral outcomes observed during opioid withdrawal. Specifically, probiotic supplementation appears to restore balance to the gut microbiome, downregulate specific serotonin receptors, and reduce anxiety-like behavior in morphine-withdrawn mice. These findings not only support the gut-brain axis as a key mediator of opioid withdrawal symptoms but also propose probiotic therapy as a promising adjunct treatment for managing the emotional and psychological challenges associated with opioid detoxification. Future studies should explore the clinical implications of these results, particularly in the context of human opioid withdrawal, to determine the feasibility of implementing probiotic-based therapies in opioid recovery programs.

The authors report there are no competing interests to declare.

This work was supported by the National Institute on Drug Abuse, National Institute of Health, Bethesda, MD under Grants: R01 DA050542, R01 DA047089, R01 DA044582, R01 DA043252, R01 DA037843, R01 DA034582, T32 DA045734, FDOH 20B12.

The data that support the findings of this study are openly available in the Harvard Dataverse at https://dataverse.harvard.edu/dataverse/GMBANX (https://doi.org/10.7910/DVN/ZXEWRE, https://doi.org/10.7910/DVN/YCP2HH, https://doi.org/10.7910/DVN/NL1Y3I, https://doi.org/10.7910/DVN/5XJKQC, https://doi.org/10.7910/DVN/RLUNWT, https://doi.org/10.7910/DVN/K07FX0)

The generative AI ChatGPT-4-turbo was employed to improve grammar and flow of introduction and discussion sections.

The authors thank Eridania Valdes for technical assistance as well as Dr. Eleonore Beurel for advice on experimental design and for use of equipment.

