## Supplementary figures and images for "Anxiety-like behavior during protracted morphine withdrawal is driven by gut microbial dysbiosis and attenuated with probiotic treatment"

### Supplemental figures

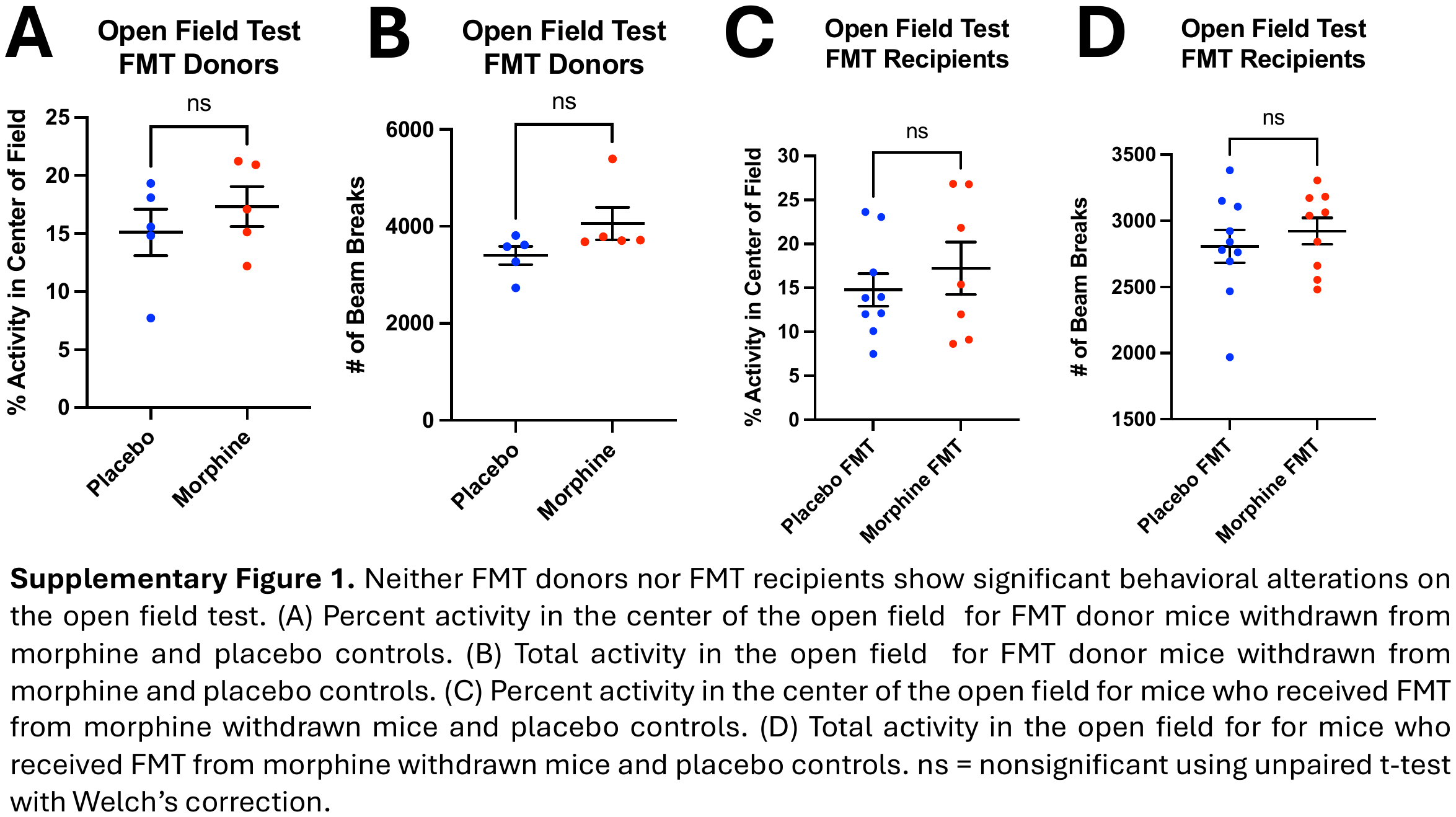

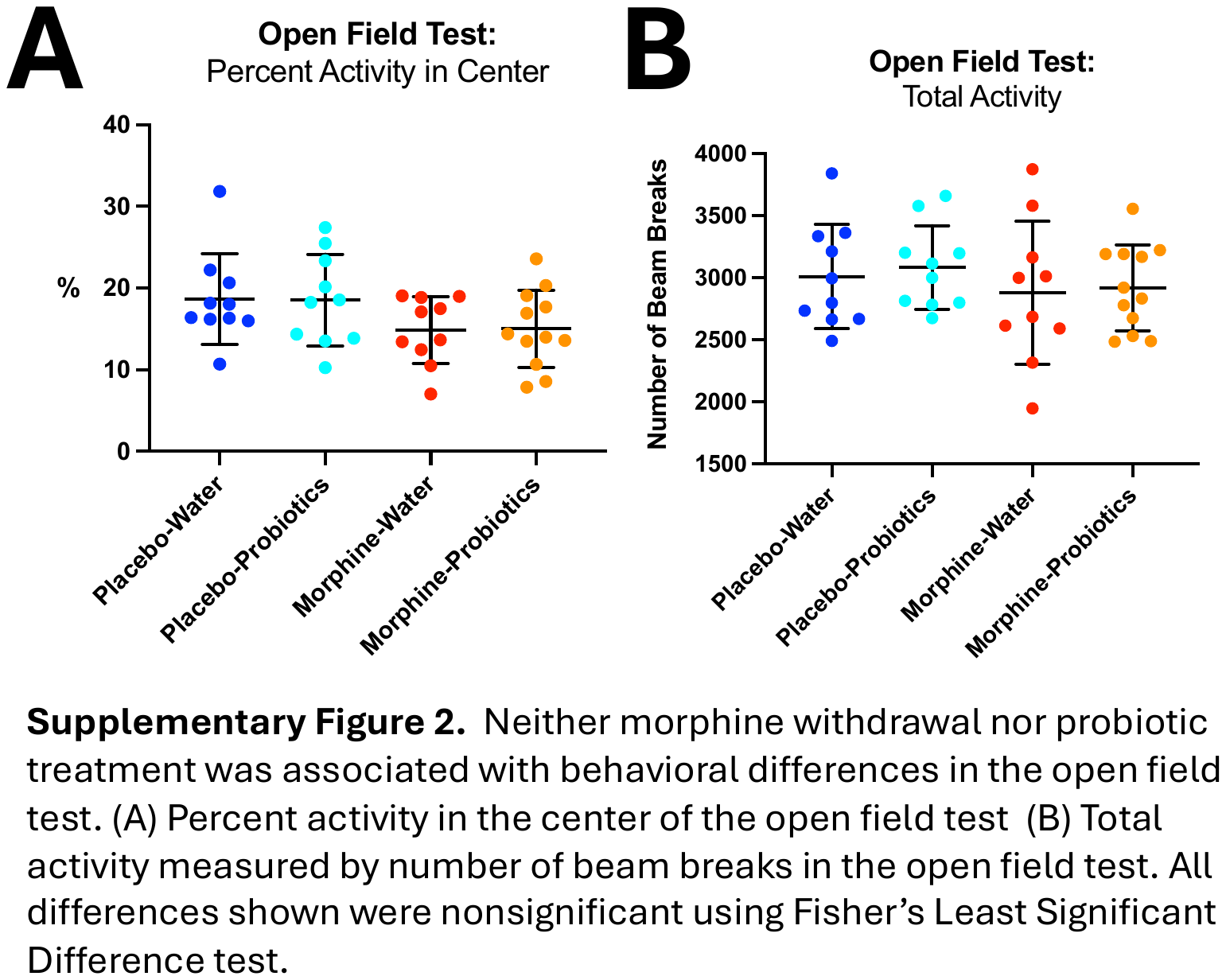
